# Serological surveillance for wild rodent infection with SARS-CoV-2 in Europe

**DOI:** 10.1101/2022.08.02.502439

**Authors:** Vincent Bourret, Lara Dutra, Hussein Alburkat, Sanna Mäki, Ella Lintunen, Marine Wasniewski, Ravi Kant, Maciej Grzybek, Vinaya Venkat, Hayder Asad, Julien Pradel, Marie Bouilloud, Herwig Leirs, Valeria Carolina Colombo, Vincent Sluydts, Peter Stuart, Andrew McManus, Jana A. Eccard, Jasmin Firozpoor, Christian Imholt, Joanna Nowicka, Aleksander Goll, Nathan Ranc, Guillaume Castel, Nathalie Charbonnel, Tarja Sironen

**Affiliations:** University of Helsinki: Medicum, 00014 Helsinki and Veterinary Medicine, 00790 Helsinki, Finland; INRAE - Université de Toulouse UR 0035 CEFS, 31326 Castanet Tolosan, France; ANSES-Nancy, Laboratoire de la rage et de la faune sauvage, 54220 Malzéville, France; Department of Tropical Parasitology, Institute of Maritime and Tropical Medicine, Medical University of Gdansk, 81-519 Gdynia, Poland; CBGP, INRAE, CIRAD, IRD, Institut Agro, University of Montpellier, Montpellier, France; University of Antwerp, Evolutionary Ecology Group, 2610 Wilrijk, Belgium; Consejo Nacional de Investigaciones Científicas y Técnicas (CONICET), Buenos Aires, Argentina; Munster Technological University, Department of Biological and Pharmaceutical Sciences, Clash, V92 CX88 Tralee, Ireland; University of Potsdam, Institute of Biochemistry and Biology, 14469 Potsdam, Germany; Julius Kühn Institute, 48161 Münster, Germany

**Keywords:** Europe, rodents, SARS-CoV-2, serology

## Abstract

We report serological surveillance for exposure to SARS-CoV-2 in 1,237 wild rodents and other small mammals across Europe. All samples were negative with the possible exception of one. Given the ongoing circulation of this virus in humans and potential host jumps, we suggest such surveillance be continued.

## Main text

Reverse zoonotic disease transmission of diverse pathogens (bacteria, viruses, eukaryotic parasites, fungi) from humans to animals has been recognized as a major concern for years and documented globally (1). When such human pathogens become established in an animal population, this population may act as a reservoir for human re-infection, hampering or preventing pathogen eradication. Circulation in new animal hosts may also lead mutation-prone pathogens, such as RNA viruses, to accumulate new mutations (2), with potential for unforeseen consequences for human epidemiology (3). Clearly, in the context of the COVID-19 pandemic and from a One Health perspective, “there is an urgent need to develop frameworks to assess the risk of SARS-CoV-2 becoming established in wild mammal populations” (4). On March 7^th^, 2022, the FAO, OIE and WHO issued a joint statement to “promote monitoring of wildlife and encourage sampling of wild animals known to be potentially susceptible to SARS-CoV-2”, and “emphasize the importance of monitoring mammalian wildlife populations for SARS-CoV-2 infection”. Wild rodents in particular have been suggested to be among the more susceptible species to SARS-CoV-2 infection, and a range of rodent species have confirmed susceptibility to experimental infection (5–7). The exact course of infection may differ between rodent host species, but it generally results in little or no detectable disease, infectious virus shedding for 4-7 days post infection and transmission to naïve contact individuals (5–7). These characteristics suggest a clear potential for reverse transmission, broad circulation and possible long-term establishment in rodent populations. Such an event would certainly be of concern - hamsters, for instance, have transmitted this virus back to humans with onward human-to-human transmission (8). Consequently, on December 6^th^, 2021 the FAO-OIE Advisory Group on SARS-CoV-2 Evolution in Animals indicated that the lack of a large surveillance study of rodent populations exposed to human contact was a major gap in SARS-CoV-2 research.

Animal experiments have shown that antibodies are consistently detected for at least several weeks after rodent infection with SARS-CoV-2 (5–7). When field prevalence is low or unknown, serology is therefore the method of choice to maximize the chances of detecting the circulation of such viruses that cause brief infection but a longer-lasting serological response. A recent survey found a sewage rat (*Rattus* spp.) in Hong Kong to be seropositive for SARS-CoV-2 (9), and considering the high biodiversity and ubiquity of rodents, broader surveillance studies in other continents, habitats, and non-commensal rodent species are urgently needed.

To shed light on the possible transmission and establishment of SARS-CoV-2 in wild rodents in different settings, we conducted a large-scale serological survey of SARS-CoV-2 in multiple rodent species across Europe. We surveyed urban parks (including zoos) which offer ample opportunity for transmission between humans and rodents, and forests, since other wild forest mammals have naturally become infected with SARS-CoV-2. We sampled 1,202 rodents and 35 Soricidæ (*Sorex* and *Crocidura*) from 23 forests sites and 8 urban parks in five European countries (Ireland, Belgium, France, Germany and Poland) during 2021 (Figure 1 and Supplementary file: Table S1). We then assessed each individual’s SARS-CoV-2 serological status using an infected cell-based immunofluorescent (IF) assay (10) (see assay details in the Appendix).

**Figure 1.**
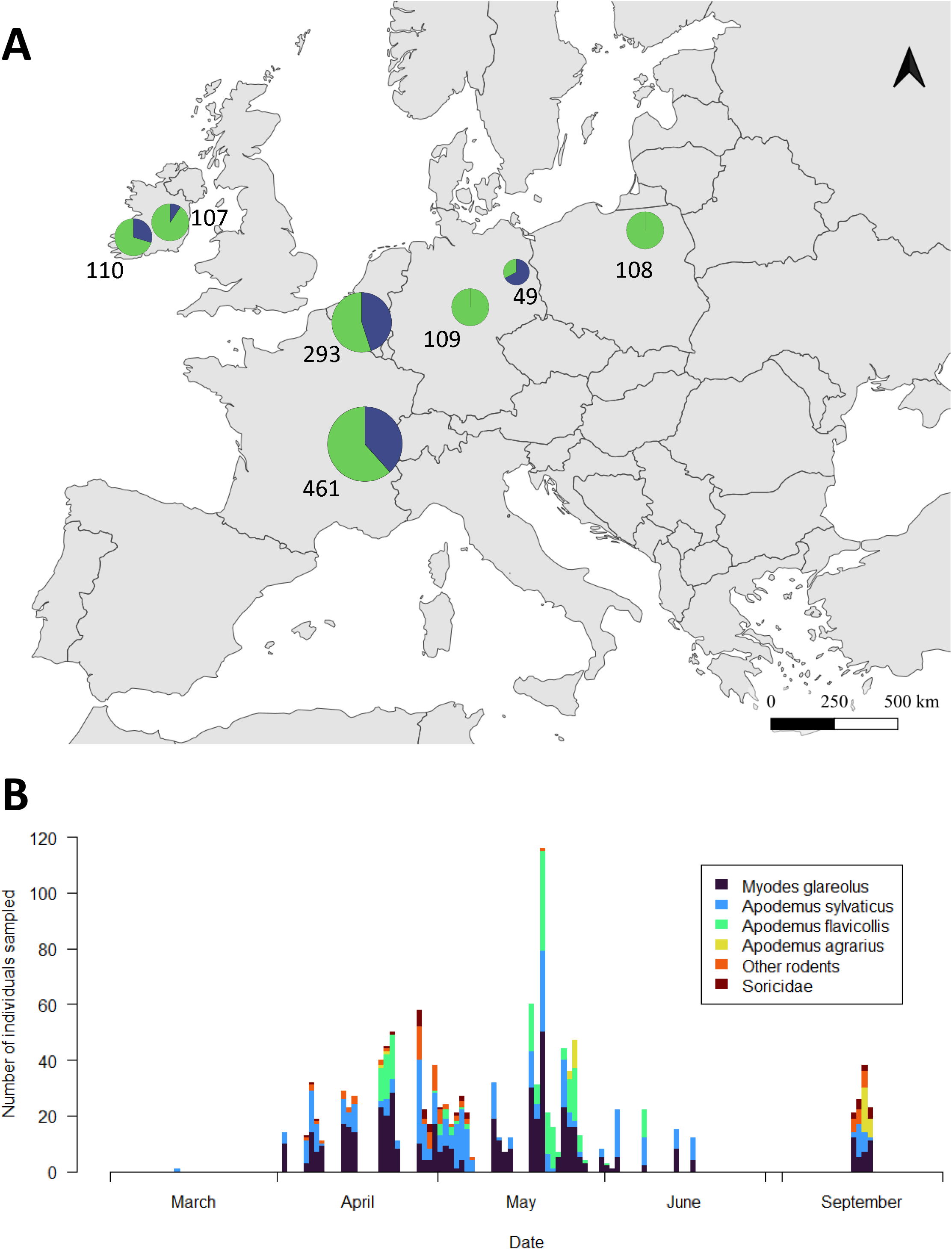
Sampling of various European areas to detect SARS-CoV-2 antibody response in wild rodents. **A**, location of sampling areas. Colours indicate the proportion of samples taken in the two habitat types (green: forests; blue: urban parks) and symbol size and numbers indicate sample size. Samples were taken from up to eight different sites in each country (detail of sampling periods, habitats and rodent species is in Supplementary Table S1). **B**, number of individuals sampled by date and taxonomy.

All but one of the rodents sampled were seronegative for SARS-CoV-2. The one suspected positive individual (assayed twice by IF and both times unambiguously positive, see representative images in Appendix) was a wood mouse (*Apodemus sylvaticus*) sampled in an urban park near the city of Antwerp, Belgium on April 6^th^ 2021. To further investigate potential virus circulation in the area, we carried out a SARS-CoV-2 specific PCR on samples from all 59 individuals from that particular location (methods details in the Appendix). These PCR were all negative including the seropositive individual. This could be expected, given the short virus shedding period described in rodents (5–7).

On July 6^th^, 2022, the OIE pointed out that “While occasional occurrences of COVID-19 in domestic or zoo animals show little long-term consequence, infections at wildlife population levels indicates the possibility of further evolution of the virus in animals, and a future reintroduction of the virus into humans at a later date. A worrying possibility.” The agency also re-stated that “Only through monitoring the virus’s reach can we understand the full picture of animal and human health, and effectively predict and prevent future outbreaks of the disease.” This survey shows that SARS-CoV-2 had not spread to a high prevalence in Northern European wild rodents as of April-September 2021. Nonetheless, the current circulation in humans of new variants with potential rodent-tropic features “is a stark warning of the risk of reservoirs of SARS-CoV-2 in wild rodents” (3), and calls for continued SARS-CoV-2 surveillance in wild rodents.

## Supporting information

images in Appendix

## Acknowledgements

We are most grateful to Jussi Hepojoki at Medicum, University of Helsinki for information and advice on the IF assay. We are also grateful to various staff at the Dept. of Veterinary Medicine at Helsinki University: Sofia Greilich and Akseli Valta who helped prepare IFA slides, and Maija Suvanto and Ruut Uusitalo who helped set up the RNA extraction protocol. We thank the animal experiment team at ANSES LRFSN for animal care and sample collection, and Jens Jacob from the Julius Kühn Institute for supporting the project in Germany. Sampling from Thuringia (Germany) was funded by the DFG Priority Program 1374 “Biodiversity - Exploratory”, and we thank the fieldwork organisers, local management teams, data management team, the Hainich national park administration and the land owners for collaboration.

This research was funded through the European H2020 (WP 2018-2020) call and the 2018–2019 BiodivERsA joint call for research proposals, under the BiodivErsA3 ERA-Net COFUND programme, and co-funded by Agence Nationale de la Recherche (ANR), Research Foundation–Flanders (FWO), National Science Centre, Poland (NCN), Deutsche Forschungsgemeinschaft (DFG) and the EPA Research Programme 2021-2030. The EPA Research Programme is a Government of Ireland initiative funded by the Department of the Environment, Climate and Communications. MG, JN and AG were supported by the National Science Centre, Poland, under the BiodivERsA3 programme (2019/31/Z/NZ8/04028).

The authors declare that there are no conflicts of interests.

## Biographical sketch

Vincent Bourret is a D.V.M with a Ph.D in virology from University of Cambridge, UK. He is now a researcher at INRAE, France, and works on wildlife disease ecology.

