## Appendix for "Serological surveillance for wild rodent infection with SARS-CoV-2 in Europe"

by V. Bourret *et al.* 2022

This Appendix contains methodological details on sample collection (field samples and experimental controls) and the laboratory diagnostic assays (serological and molecular.) The final section consists of the work's legal and ethical statements.

### A. Sampling

#### *1) Field samples*

Small mammals were trapped using snap-traps (Germany) or live traps (all countries) whose size was adapted to the target species (e.g. large wire mesh traps for rats, Sherman traps for *Myodes*, *Microtus*, or *Apodemus* sp). Traps were set either following predetermined transect lines, or at specific sites where rodents had been seen by site managers (details available upon request.) Traps were left on each sampling site for one to eleven nights. Live traps were checked every morning, and re-filled with hydrophobic cotton and food (weeds, carrots, sardine, peanut butter) daily to provide resources and a suitable environment for trapped animals.

Trapped animals were identified using morphological criteria in the field. When field identification was problematic, molecular identification was performed in the lab: *Microtus* species were identified using Sanger sequencing of CO1 fragment (1) and *Apodemus sylvaticus* and *A. flavicollis* were distinguished using the AP-PCR as in (2).

Live-trapped animals were euthanized with an overdose of isoflurane. Rodents found dead by site managers were also included in the study. All animals were dissected and several organs were collected (including the heart, placed in PBS for serological assaying, and colon, placed in RNAlater), and stored at -20 °C until assayed. Individual characteristics were also recorded (mass, body length, gender, sexual characteristics.)

A breakdown of the samples collected by host species, localities and dates is provided in Supplementary file Table S1.

All legal and ethical information regarding this study is collated in a dedicated section at the end of this Appendix.

#### *2) Experimental control samples*

##### *a. Vaccinated animals*

Ten-week old Syrian golden hamsters (*Mesocricetus auratus*) were acclimatized at the University of Helsinki biosafety level 3 (BSL-3) facility for seven days in individually ventilated biocontainment cages (ISOcage; Scanbur) with one hamster per cage. Animals were then immunized twice seven days apart with an experimental receptor binding domain-based nasal vaccine (patent pending.) Immunized hamsters were euthanized by cervical dislocation 14 days after the second immunization and heart was collected into PBS and stored at -20 °C.

b. *Challenged animals*

***Virus.*** The challenge SARS-CoV-2 strain used in these experiments was prepared as described previously (3) and used at passage 2.

***Animals.*** Seven week-old female Syrian golden hamsters (strain RjHan:AURA) were purchased from Janviers's breeding Centre (Le Genest, St Isle, France), housed in an animal-biosafety level 3 facility at ANSES, Malzéville, France and left to acclimatize for a minimum of seven days before challenge. For collection of both sample types (below), the animals were anesthetized with a mix of ketamine + xylazine (150 mg/kg + 10 mg/kg) administered by the intraperitoneal route, euthanized by exsanguination, and necropsied.

***Plasma samples.*** Six hamsters were anesthetized using isoflurane and intranasally inoculated with 40 µL containing $10^4$ $TCID_{50}$ of SARS-CoV-2 virus (20 µL in each nostril). At fourteen days post-infection, the blood was collected by heart puncture in 4 mL EDTA 3K Vacutest tubes. The plasmas were obtained after centrifugation (15 min, 1000 g) and stored at -16 °C until analysis. The presence of SARS-CoV-2 neutralizing antibodies was confirmed in these samples by seroneutralisation prior to IFA testing.

***Heart samples.*** Six hamsters were anesthetized with isoflurane and intranasally inoculated with 40 µL containing $10^5$ $TCID_{50}$ of virus (20 µL in each nostril). At fifteen days post-infection, after exsanguination, the hearts were collected in vials containing 500 µL of sterile PBS. The samples were then stored at -16 °C until analysis. The presence of SARS-CoV-2 antibodies in these samples was confirmed by microsphere immunoassay prior to IFA testing.

Figure S1 shows representative IFA slides, including three different positive controls and three field samples (two negative and the one positive.)

### B. Laboratory diagnostic procedures

#### 1) *Immunofluorescent assay (IFA)*

All field samples were screened for SARS-CoV-2 antibodies using the immunofluorescent assay (IFA) described in (4) with the following modifications:

(i) samples consisted of whole rodent hearts in PBS, whose supernatant was assayed undiluted, and

(ii) the secondary antibody was fluorescein isothiocyanate (FITC)-conjugated anti-mouse IgG.

The capacity of this test for robust detection of rodent SARS-CoV-2 antibody response was assessed using a range of animal experiments comparing:

- different immunization methods (vaccination vs. experimental infection),
- different sample types (plasma vs. heart in PBS),
- different secondary conjugates (anti-mouse vs. anti-hamster),

as presented in Table S2. Results were positive for all tested combinations of the above factors (Table S2.) The corresponding animal procedures are detailed in section A.2. above.

**Table S2.** Characteristics of hamster positive controls used to assess the ability of the immunofluorescent assay to detect rodents seropositive for SARS-CoV-2.

| Sample name | Immunization method | Days post immunization | Sample type | Secundary conjugate | IFA result |
|---|---|---|---|---|---|
| pos176 | Vaccination | 14 | Heart in PBS | Anti-mouse | + |
| pos177 | Vaccination | 14 | Heart in PBS | Anti-mouse | + |
| pos178 | Vaccination | 14 | Heart in PBS | Anti-mouse | + |
| pos176 | Vaccination | 14 | Heart in PBS | Anti-hamster | + |
| pos177 | Vaccination | 14 | Heart in PBS | Anti-hamster | + |
| pos178 | Vaccination | 14 | Heart in PBS | Anti-hamster | + |
| H1242 | Infection | 14 | Plasma | Anti-mouse | + |
| H1243 | Infection | 14 | Plasma | Anti-mouse | + |
| H1244 | Infection | 14 | Plasma | Anti-mouse | + |
| H1245 | Infection | 14 | Plasma | Anti-mouse | + |
| H1246 | Infection | 14 | Plasma | Anti-mouse | + |
| H1247 | Infection | 14 | Plasma | Anti-mouse | + |
| H1242 | Infection | 14 | Plasma | Anti-hamster | + |
| H1243 | Infection | 14 | Plasma | Anti-hamster | + |
| H1244 | Infection | 14 | Plasma | Anti-hamster | + |
| H1245 | Infection | 14 | Plasma | Anti-hamster | + |
| H1246 | Infection | 14 | Plasma | Anti-hamster | + |
| H1247 | Infection | 14 | Plasma | Anti-hamster | + |
| H4891 | Infection | 15 | Heart in PBS | Anti-mouse | + |
| H4892 | Infection | 15 | Heart in PBS | Anti-mouse | + |
| H4893 | Infection | 15 | Heart in PBS | Anti-mouse | + |
| H4894 | Infection | 15 | Heart in PBS | Anti-mouse | + |
| H4895 | Infection | 15 | Heart in PBS | Anti-mouse | + |
| H4896 | Infection | 15 | Heart in PBS | Anti-mouse | + |

#### *2) PCR screening of Fort 6 samples (Belgium)*

The 59 colon samples from the Fort 6 location near Antwerp, Belgium, where the seropositive individual had been detected, were screened for SARS-CoV-2 infection using a specific PCR. Total RNA was extracted from rodent tissue samples using the QIAamp 96 Virus QIAcube HT kit (Qiagen). Colon samples were first removed from RNAlater in a BSL2 laboratory and approx. 10 mg were placed in 180 µL ATL buffer + 20 µL proteinase K (supplied with the kit) in secure 2 mL tubes containing two glass bead and autoclaved sand. The tubes were then incubated at 56 °C for 30 min for enzymatic lysis, after which they were shaken at 30 Hz for 2 x 2 min using a TissueLyser (Qiagen.) Lysates were then spun at 500 g for 5 minutes, and 200 µL clear supernatant were used as starting material for automated QIAcube extraction as per manufacturer's instruction, with the following modification: the final target elution volume was 120 µL. Eluted RNA were then stored at -80 °C until assayed by PCR. Tissues from SARS-CoV-2 challenged rodents were used as positive extractions controls with every extraction batch and returned consistent positive PCR.

Extracted RNA were then assayed using the Luna SARS-CoV-2 RT-qPCR Multiplex Assay Kit (New England BioLabs Inc, MA USA) as per manufacturer's instructions with no modification (using the provided kit positive control.)

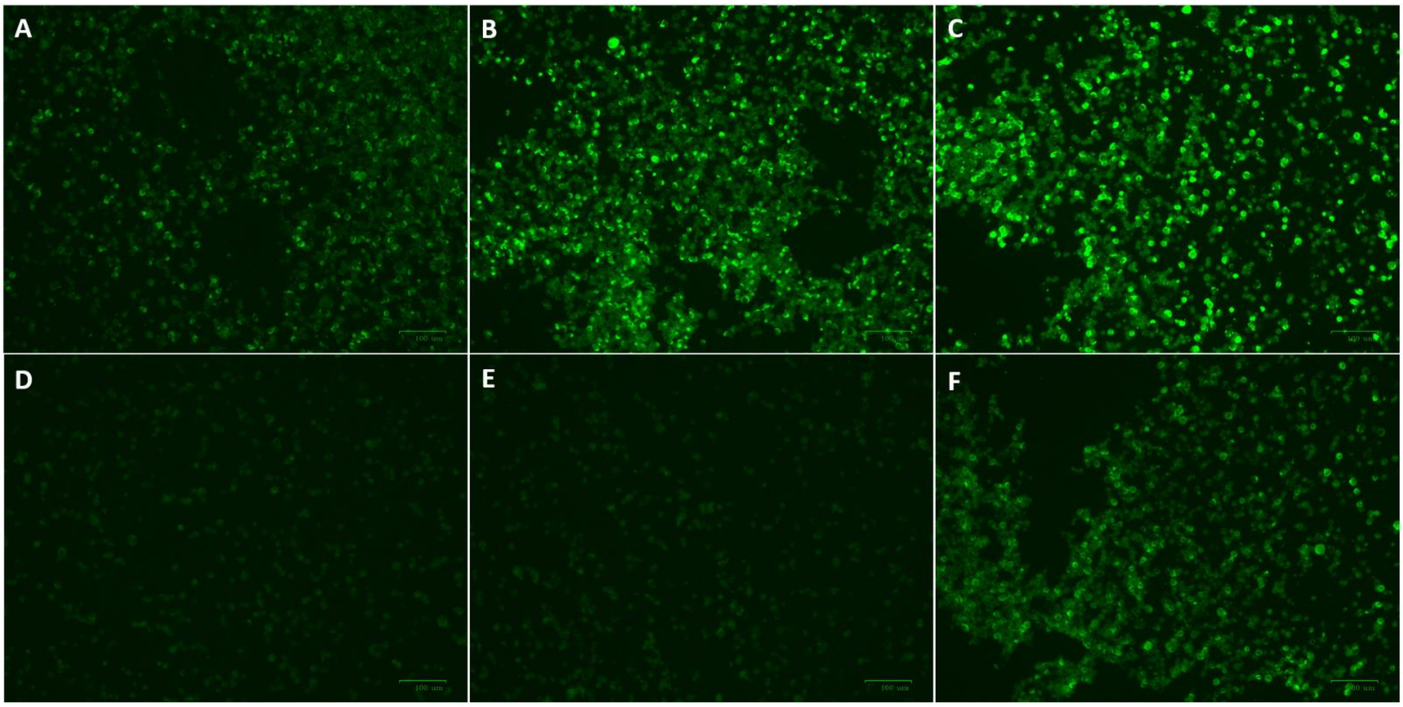


**Figure S1.** Representative images of SARS-CoV-2 immunofluorescent assays (IFA) in rodents.

***Top row: positive controls*** (*A:* vaccinated hamster, heart in PBS. *B:* challenged hamster, plasma, anti-mouse secondary conjugate. *C:* challenged hamster, plasma, anti-hamster secondary conjugate.)

***Bottom row: field samples*** (*D:* negative *Myodes glareolus*; *E:* negative *Apodemus sylvaticus*; *F:* the positive *Apodemus sylvaticus*.)

Scale bar at the bottom right of each photograph represents 100 µm. Note the presence of both positive (brighter green) and negative (dim) cells in comparable proportions in positive tests (A, B, C, F), while the negative tests show exclusively negative cells (D, E.)

### C. Legal and ethical statements

#### 1) *Field sampling*

*a. Belgium*

The procedures were approved by the University of Antwerp Ethical Committee for Animal Experiments (permit number 2020-21). Small mammal trapping was approved by the Flemish regional nature authority (Agentschap voor Natuur en Bos, ANB). The handling of small mammals was carried out in accordance with the recommendations in Directive 2010/63/EU.

*b. France*

The procedures complied with the French regulations on care and protection of laboratory animals (French Law 2001-486 issued on June 6th, 2001 and Directive 2010/63/EU issued on September 22nd, 2010.) They were also authorised by the regional ethical committee for animal experiments (Languedoc Roussillon, n°36, 2020-2025.) The CBGP laboratory, which carried out the sampling in France, has approval (no. D-34-169-003) from the Departmental Direction of Population Protection (DDPP, Hérault, France) for the sampling of rodents and the storage and use of their tissues.

*c. Germany*

The study was performed in accordance with the applicable international and institutional guidelines for the use of animals in research. In Brandenburg, collection of rodents was performed under the permission of “Landesamt für Arbeitsschutz, Verbraucherschutz und Gesundheit Brandenburg (LAVG)" (no. 2347-A-16-1-2020, for procedure) and “Landesamt für Umwelt Brandenburg (LfU)" (no. LFU-N1-4744/97+17#194297/2020, for sites and species exemptions.) In Thüringen, all procedures were permitted by the "Thüringer Landesamt für Verbraucherschutz (TLV)" (no. 22-2684-04-15-105/16.)

*d. Ireland*

Ethical approval for the work carried out in Ireland was obtained from the Institute of Technology (Tralee, Ireland) Research Ethics Committee, and following that the Health Products Regulatory Authority (HPRA) granted authorisation (Authorisation Number: AE22171/I004) for euthanasia of the rodents to be sampled.

*e. Poland*

This study was carried out according to the recommendations in the Guidelines for the Care and Use of Laboratory Animals of the Polish National Ethics Committee for Animal Experimentation.

### *2) Animal experiments*

The hamster vaccination experiment (carried out in Helsinki, Finland) was approved by the Animal Experiment Board of Finland (license number: ESAVI/28687/2020.)

The SARS-COV-2 challenge experiments (carried out in Malzéville, France) complied with the 2010/63/CE regulation of the European Parliament and of the council of 22 September 2010 on the protection of animals used for scientific purposes. The experiments were approved by the Anses/ENVA/UPEC ethics committee and the French Ministry of Research (license numbers: APAFIS #32431-2021071514369893 v2 for the plasma collection experiment and APAFIS #33544-2021102114466426 v2 for the heart collection experiment.)
