## Supplementary figures and images for "Serological surveillance for wild rodent infection with SARS-CoV-2 in Europe"

### images in Appendix

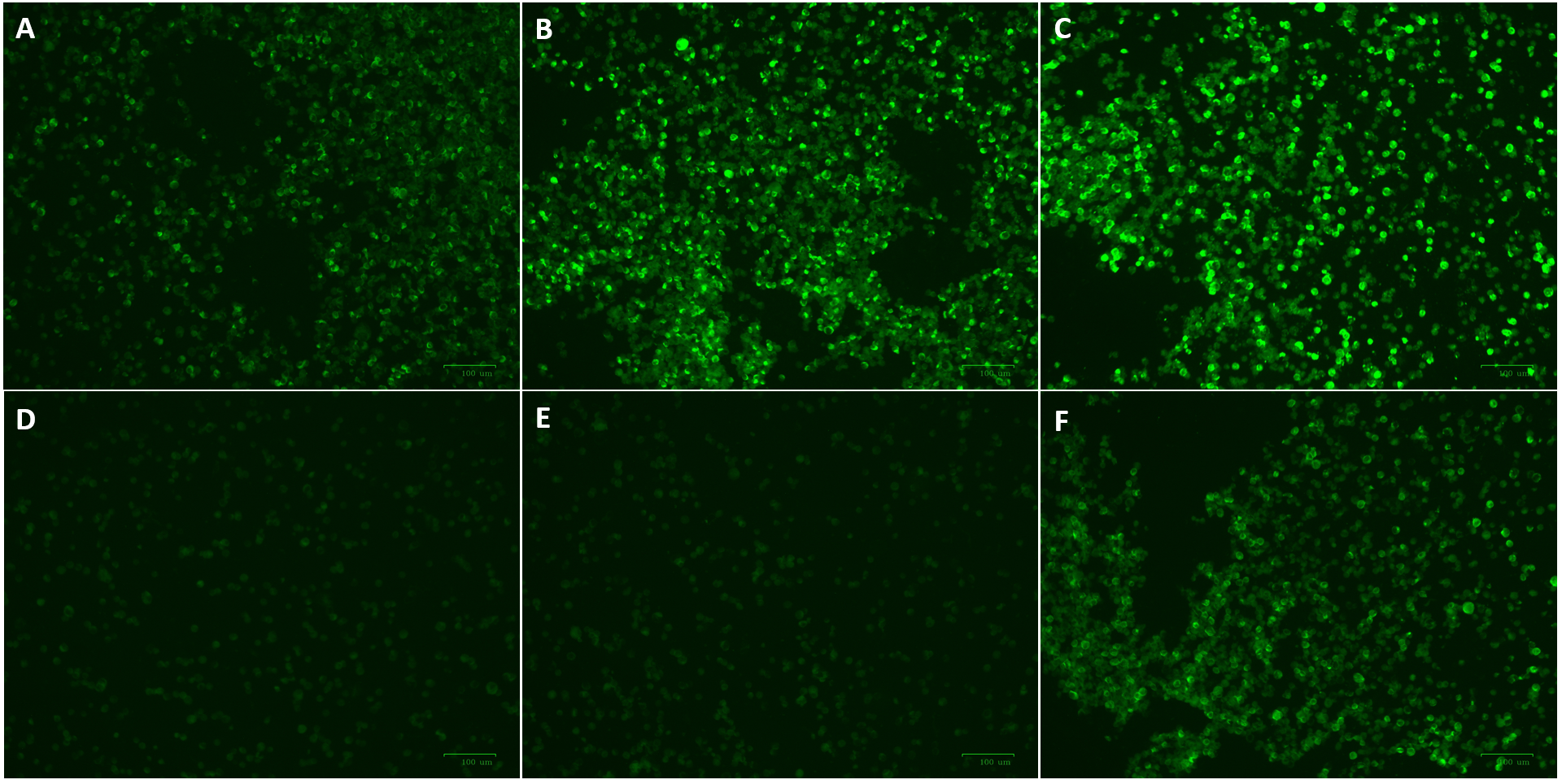
